# Substrate stiffness modulates phenotype-dependent fibroblast contractility and migration independent of TGF-β stimulation

**DOI:** 10.1101/2025.06.16.659915

**Authors:** Mirko D’Urso, Pim van den Bersselaar, Sarah Pragnere, Paolo Maiuri, Carlijn V.C. Bouten, Nicholas A. Kurniawan

**Affiliations:** Department of Biomedical Engineering, Eindhoven University of Technology, Eindhoven, The Netherlands; Institute for Complex Molecular Systems, Eindhoven University of Technology, Eindhoven, The Netherlands; Mines Saint-Etienne, Université Jean Monnet Saint-Etienne, INSERM, SAINBIOSE U1059, Saint-Etienne, France; Department of Molecular Medicine and Medical Biotechnology, Università degli Studi di Napoli “Federico II”, Naples, Italy

## Abstract

During wound healing, fibroblasts undergo radical processes that impact their phenotype and behavior. They are activated, recruited to the injury site, assume a contractile phenotype, and secrete extracellular matrix proteins to orchestrate tissue repair. Thus, fibroblasts response require dynamic changes in cytoskeleton assembly and organization, adhesion morphology, and force generation. At the same time, fibroblasts experience changes in environmental stiffness during tissue wounding and healing. Although cells are generally known to use their adhesion–contraction machinery to sense microenvironmental stiffness, little is known about how stiffness affects the fibroblast phenotypical transition and behavior in wound healing. Here we demonstrate that stiffness plays a deterministic role in determining fibroblast phenotype, surprisingly even overruling the classical TGF-β-mediated stimulation. By combining morphometric analysis, traction force microscopy, and single-cell migration analysis, we show that environmental stiffness primes the cytoskeletal and mechanical responses of fibroblasts, strongly modulating their morphology, force generation, and migration behavior. Our study, therefore, points to the importance of tissue stiffness as a key mechanobiological regulator of fibroblast behavior, thus serving as a potential target for controlling tissue repair.

## Introduction

Fibroblasts are a highly heterogeneous cell type involved in regulating tissue homeostasis by maintaining tissue microarchitecture. They are found in almost every human tissue. Despite the tissue-specific properties unique to each fibroblast lineage, all fibroblasts share the ability to modify the microenvironment through extracellular matrix (ECM) production and remodeling[1,2]. Importantly, when tissue microarchitecture homeostasis is disrupted, for example due to wounding, the composition, density, and organization of the ECM are altered, often resulting in overall tissue stiffening that can evoke fibroblast response[3,4]. Moreover, upon initiation of wound-associated immune response, fibroblasts become activated and increase their production of ECM components, which again leads to tissue stiffening[5]. If not regulated properly, this positive feedback loop and overproduction of ECM will eventually lead to pathological conditions such as fibrosis[6–8].

Concurrent with tissue wounding and the subsequent acute inflammatory response, an immediate and local increase in the concentration of TGF-β[9,10], which induces fibroblast activation and initiates the fibroblast-to-myofibroblast transition (FMT). As the insult to the tissue resolves myofibroblasts generally undergo apoptosis. However, under chronic pro-inflammatory conditions, myofibroblast apoptosis does not take place[11–13]. As a result, the presence of cells with persistent myofibroblastic phenotype is a well-known hallmark of pathological conditions, particularly in fibrotic tissues[13–15].

There is a growing body of evidence that fibroblast phenotype is modulated by mechanical forces at several regulatory levels. First, fibroblasts and myofibroblasts produce and release TGF-β in a closed, inactive conformation, which is then embedded in the ECM as a large latent complex[16]. When cells exert force on the ECM, TGF-β changes its conformation, allowing it to bind to the membrane receptors of fibroblasts, leading to a positive loop whereby FMT is induced and more TGF-β is produced[16]. Second, in their activated state, fibroblasts migrate towards the site of tissue disruption, facilitated by an elevated secretion of matrix metalloproteinases for ECM degradation. Cell migration requires active force generation and cytoskeletal rearrangements that influence the mechanical properties of both the cells and the ECM[17–20]. At the same time, the phenotypic shift to a myofibroblastic state strongly increases cell contractility, as enabled by the inclusion of alpha-smooth muscle actin (αSMA) within the cytoskeleton, particularly in the actin filament bundles[21]. This increased contractility, in turn, plays an essential role in defining the typical spread-out morphological features of myofibroblasts and in modifying the microarchitecture of the ECM, which results in tissue stiffening[22–24].

It is noteworthy that, at the cell level, the same set of intracellular machinery is responsible for different cellular processes such as phenotypical shift, migration, and mechanosensing. Cell forces are generated by the actomyosin stress fibers and transmitted to the ECM via focal adhesions (FAs). The actomyosin stress fibers are critical for sensing microenvironmental cues and facilitating cell migration[25] whereas FAs enable cells to probe the fibroblast’s physical environment[26]. It has been shown that physical cues, such as contact guidance, can change fibroblast behavior by affecting the morphology, organization, and localization of FAs[22,27,28], leading to changes in cytoskeletal organization[6,22]. Indeed, FA morphology is closely linked to cell traction forces, and traction forces are related to cell dynamics such as migration. For example, stronger traction forces are generally associated with slower cell migration[29]. This is particularly relevant for fibroblasts, as during wound healing, dermal fibroblasts need to apply forces to the ECM to migrate and contract to close the wound [11,28,30]. The FA morphology is closely linked to the cell traction forces[31] and has been implicated in fibroblast activation[22]. Moreover, it was shown that force exerted by activated fibroblasts is higher than that of quiescent fibroblasts[30]. Many aspects of this complex multifaceted regulation have been studied individually, and there is no comprehensive framework for investigating and understanding the interplay between fibroblast phenotype, migration, cell and ECM mechanics, and cell morphology, especially in the context of FMT.

It is clear that tissue stiffness is an important factor of not only FMT, but also the underlying cellular phenomena, including cytoskeletal and FA reorganization, as well as changes in fibroblast morphology and migration. However, to which extent stiffness sensing plays a role in regulating FMT is not yet fully understood. To investigate this, in this study we examined how substrate stiffness recapitulating the stiffnesses of healthy and pathological tissues regulates the transition from fibroblast to myofibroblast phenotype via mechanosensing regulation. Quantitative characterization of fibroblast migratory behavior, traction forces, and morphology revealed a direct correlation between fibroblast phenotype, morphotype, and mechanotype along the phenotype transition in FMT. This finding thus provides a holistic insight into the cellular regulation of FMT, placing the mechanical and physical (rather than biochemical, such as in the case of TGF-β) environment of fibroblasts at the forefront of wound healing physiology.

## Results

### Stiffness sensing influences FMT regulation

To study the effect of substrate stiffness on the regulation of fibroblast phenotype, we cultured fibroblasts at sub-confluent densities on glass and polyacrylamide (pAA) hydrogels and evaluated their shape together with cytoskeleton rearrangement and focal adhesions at 2 different time points for glass, 48 and 96 hours, and after 96 hours on pAA hydrogels. These time points were chosen as, after activation by TGF-β, fibroblasts are known to take about 48 hours to enter a proto-myofibroblast state^21,22^ and approximately 96 hours for a complete myofibroblastic transition on glass[21,22]. As a reference, on glass, fibroblasts showed a dense network of F-actin bundles (Figure 1A, B), as expected for adherent cells on a stiff substrates[32,33]. Moreover, no αSMA incorporation in the stress fibers was observed. The addition of 10 ng/mL TGF-β for 48 hours led to a higher expression of αSMA, but again without incorporation into the actin stress fibers (Figure 1A). After 96 hours, however, a clear incorporation of αSMA into the actin stress fibers and an increased number of myofibroblasts were observed (Figure 1B). These findings are consistent with previous reports[21,22] and reflect the expected temporal progression of the phenotype transition along the FMT[14,15,34,35]. Furthermore, we observed that fibroblast adhesion on the glass substrates varied with time. At 48 hours, the cells were predominantly elongated with limited formation of FAs, which became enhanced by addition of TGF-β. At 96 hours, TGF-β treatment on stiff substrates led to a significant increase in the cell area and amount of FAs increased. This was furthermore accompanied by incorporation of αSMA into the actin cytoskeleton, corroborating a shift to the myofibroblast phenotype.

**Figure 1.**
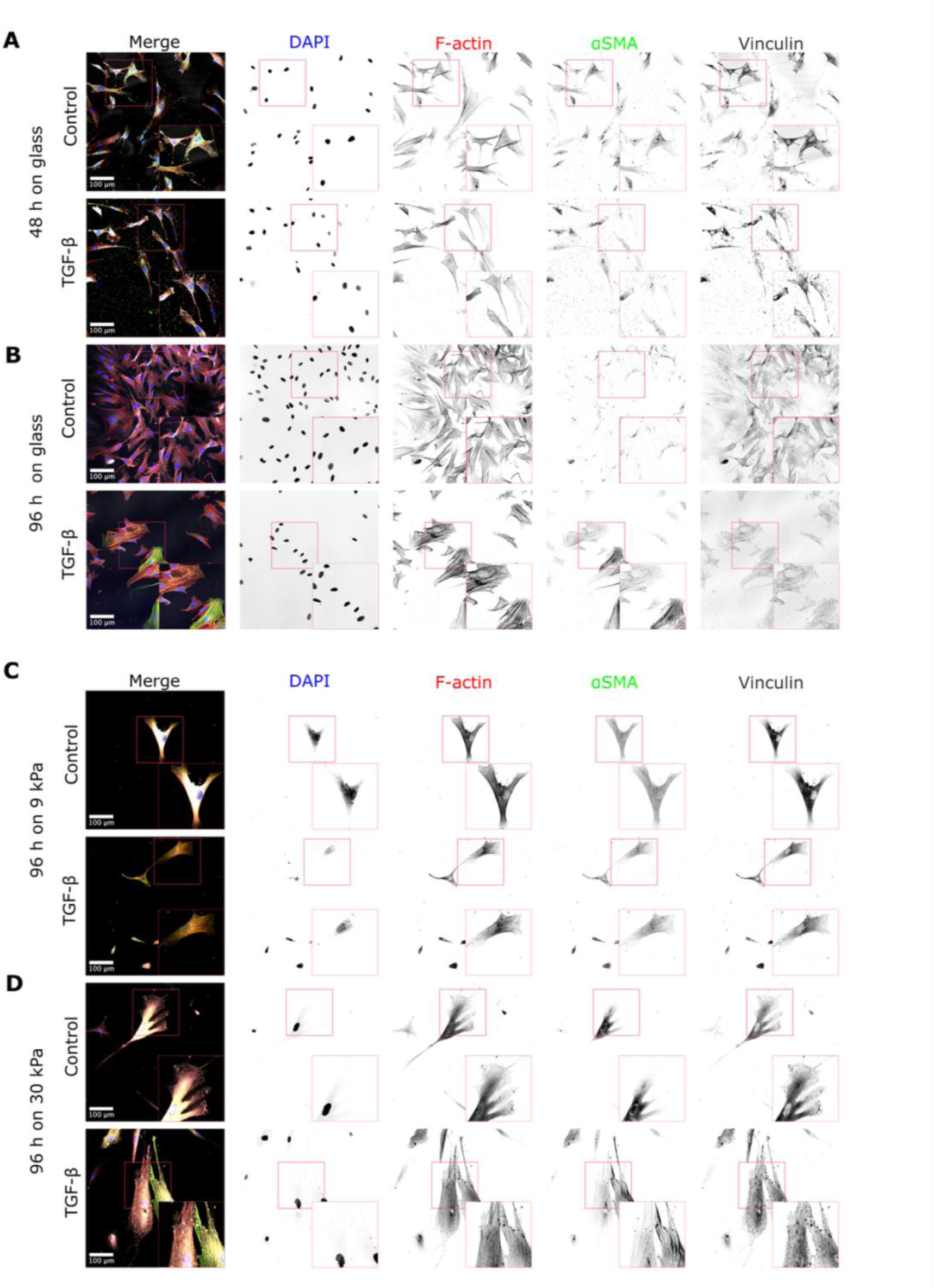
Stiffness of the substrate influences fibroblast morphology and phenotype. Fibroblasts seeded on glass for **(A)** 48 h and **(B)** 96 h with and without the addition of TGF-β. Fibroblasts seeded on pAA gels of **(C)** 9 kPa and **(D)** 30 kPa for 96 h with and without the addition of TGF-β. The merge panels show DAPI (blue), αSMA (green), F-actin (red), and vinculin (gray). Scale bar = 100 μm.

To investigate the influence of stiffness sensing in fibroblasts phenotype transition, we chose pAA gels with two stiffness values—9.3 ± 1.6 kPa and 33.3 ± 6.1 kPa (Figure S1; heretoafter referred to as ‘soft’ and ‘stiff’ gels, respectively)—to mimic those of healthy and pathological tissues[32,36,37]. By using these stiffnesses, we tested whether healthy and pathological tissue stiffnesses can be a driver for fibroblast phenotypical transition into myofibroblasts. After 96 hours on soft pAA hydrogels, fibroblasts exhibited sparse F-actin bundles with few and small focal adhesions (FAs) (Figure 1C), indicating that, despite the active actomyosin-mediated cell traction forces, the mechanical resistance from the soft substrate was not enough to support FA maturation. Interestingly, with the addition of TGF-β, we observed a more pronounced formation of actin bundles, but without an increase in cytoplasmic αSMA (Figure 1C). In contrast, on stiffer hydrogels, the actin cytoskeleton was more dense and contained more actin bundles with more numerous FAs (Figure 1D). Furthermore, the addition of TGF-β induced the αSMA incorporation into the F-actin stress fibers and the actin cytoskeleton showed thick actin bundles network together with more numerous FAs (Figure 1D).

These results show that the mechanical properties of the substrates impact both the cytoskeletal organization and the phenotype of fibroblasts. Stiffer substrates led to higher number of mature FAs, a more mature actin cytoskeleton, and αSMA incorporation within the actin stress fibers. In other words, our results suggest that the phenotypical transition from fibroblasts to myofibroblasts is primarily conditioned by stiffness mechanosensing machinery. The reduced mechanical resistance offered by soft microenvironments hampers the formation of actin bundles, which then prevents incorporation of αSMA required for the complete FMT, even in the presence of strong FMT stimulant such as TGF-β. On the other hand, a stiffer microenvironment promotes the formation of a denser cytoskeletal network with stress fibers and more FAs, facilitating fibroblast activation that is further enhanced by TGF-β.

### Fibroblast migration changes during the phenotypical transition into myofibroblast

As we have shown, different substrate stiffnesses induce morphological changes as well as cytoskeletal and FA rearrangements in fibroblasts. Importantly, cytoskeletal rearrangements are also involved in the phenotypical transition into myofibroblasts, whereby different stages in the FMT are associated with different cytoskeletal organizations[22,38–40]. We therefore hypothesized that the changes in FA and cytoskeletal organizations should translate into variations in the fibroblast migratory behavior during different phenotypical stages along FMT. To test this hypothesis, we induced fibroblast activation through the addition of TGF-β on sub-confluent cultures and examined the migration behavior of the cells at 48 and 96 hours of culture on glass substrates. Images were taken every 10 minutes for 16 hours and the trajectories of tracked cell nucleus centroids (Figure 2A and 2B) were analyzed using a custom-written single-cell migration algorithm[41] to quantify the mean cell migration speed and mean square displacement (MSD).

**Figure 2.**
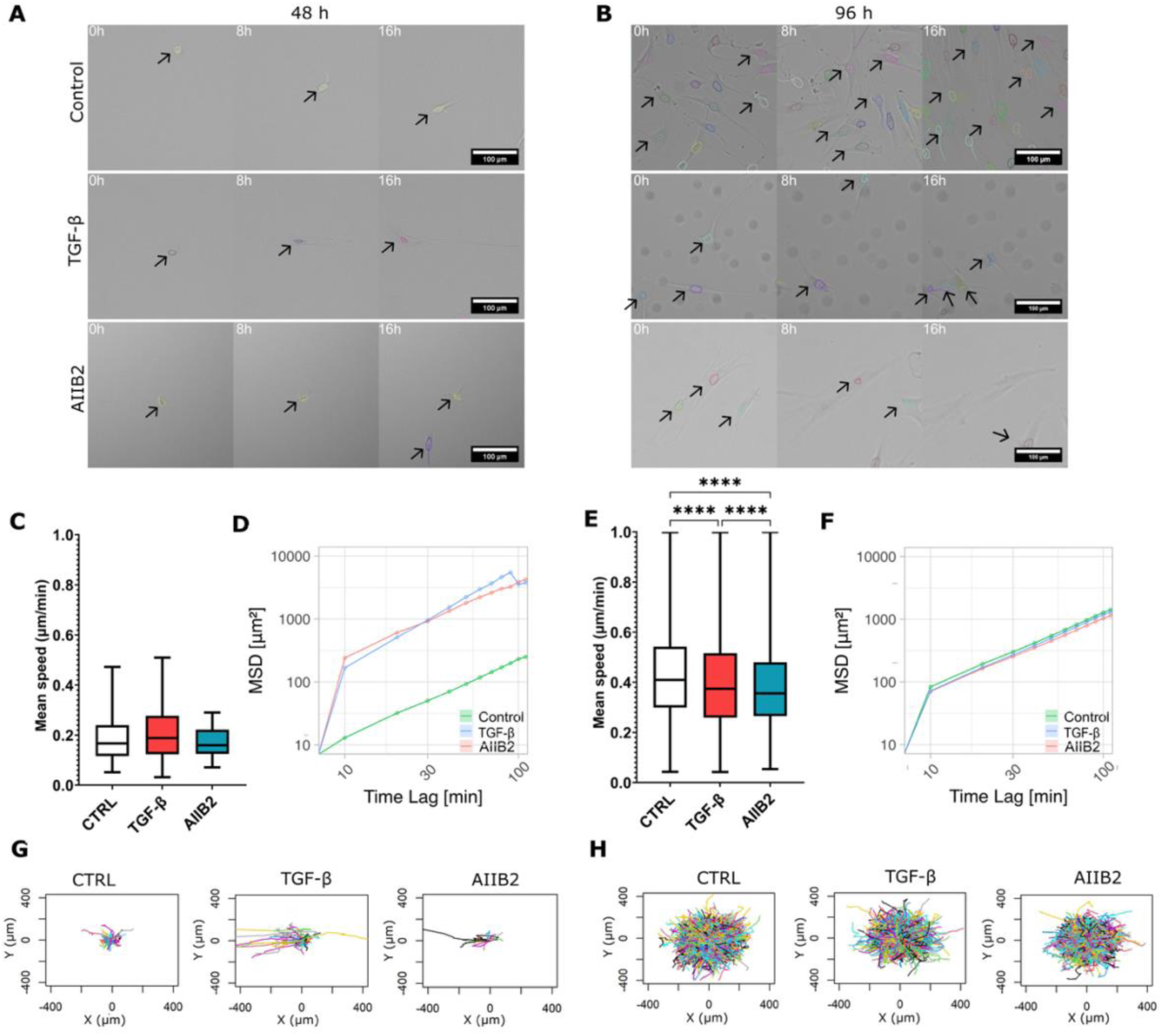
Fibroblast migration is dependent on the FMT stage. **A-B.** Representative examples of fibroblast nuclei tracking (black arrows) after (A) 48 hours and (B) 96 hours of culturing in control condition, with TGF-β stimulation, and with inhibition of β1 integrin (AIIB2). **C and E)** Mean migration speed of fibroblasts cultured in the presence and absence of TGF-β and AIIB2 after (C) 48 hours and (E) 96 hours. Data were obtained from >20 cells per condition from 6 independent samples and are shown as the boxes of the boxplots representing the quartiles of the distributions, with the whiskers indicating the outliers, ****p<0.00001 (One-way ANOVA). **D and F)** MSD of fibroblasts cultured on glass with and without TGF-β and AIIB2 for (D) 48 hours and (F) 96 hours. Data are shown as mean ± SD. **G-H)** Trajectories of fibroblasts cultured on glass with and without TGF-β and AIIB2 for (G) 48 hours and (H) 96 hours.

After 48 hours, fibroblasts treated with TGF-β were at the stage of proto-myofibroblasts, where they were activated but had not fully transitioned to be myofibroblasts[15,42]. These proto-myofibroblasts migrated with a mean speed of 0.20 ± 0.09 μm/min, with no significant difference compared to that of the fibroblasts not treated with TGF-β (0.19 ± 0.09 μm/min) (Figure 2C, *p* = 0.85). Although migration speed was not different, the MSD of the activated proto-myofibroblasts was much higher (Figure 2D), suggesting that their migration was more directed and persistent. This reflects the need for proto-myofibroblasts to quickly and persistently migrate toward a site of tissue stress *in vivo*. Indeed, the migration trajectories of these proto-myofibroblasts were more straight and persistent compared to those of unactivated fibroblasts (Figure 2G).

At 96 hours (Figure 2B), activation by TGF-β results in a complete phenotypic transition from fibroblasts to myofibroblasts (Figure 1B). Interestingly, myofibroblasts migrated with lower mean speed (0.40 ± 0.19 μm/min) compared to the control fibroblasts (0.43 ± 0.19 μm/min) (Figure 2E; *p* < 0.0001), although the MSD curves and trajectories were generally comparable between the two conditions (Figure 2F and 2H). Since myofibroblasts develop stronger substrate adhesion[30], which is important for cell migration[25,29], we hypothesized that this characteristics of reduced migration speed with higher persistence for myofibroblasts may be mediated by integrin-based focal adhesions. To test this, we introduced the inhibitor AIIB2, which blocks β1 integrin activation and prevents focal adhesion (FA) maturation (Figure S2). Consistent with our expectations, inhibition with AIIB2 led to a reduction in mean migration speed (0.39 ± 0.17 μm/min) compared to other conditions (Figure 2E; *p* < 0.0001). In contrast, inhibition with AIIB2 did not significantly affect the mean migration speed of protomyofibroblasts (0.17 ± 0.06 μm/min; Figure 2C, *p* = 0.8), nor their MSD and trajectories (Figure 2D and 2G), further suggesting that FAs play a more pronounced role in the adhesion and migration at the later stages of FMT.

This result indicates that the development of mature FAs, facilitated by incorporation of αSMA into the actin stress fibers, typifies fully transitioned myofibroblasts and strongly impacts their migratory behavior. Higher amount of mature FAs induces stronger fibroblast contractility and at the same time negatively influences cell migration. These results demonstrate the important role that FA-mediated mechanosensing plays in regulating the phenotype-dependent migratory behavior of fibroblasts.

### Cell traction forces depend on substrate stiffness and fibroblast activation

The results presented in the previous sections suggest that the distinct migration behaviors of different fibroblast phenotypes are linked to their FA-mediated contractile forces. To unravel the mechanics underlying the regulation of migratory behavior on substrates with varying stiffness, we employed traction force microscopy to measure the forces exerted by fibroblasts on the polyacrylamide (pAA) gel substrates (Figure 3A).

**Figure 3.**
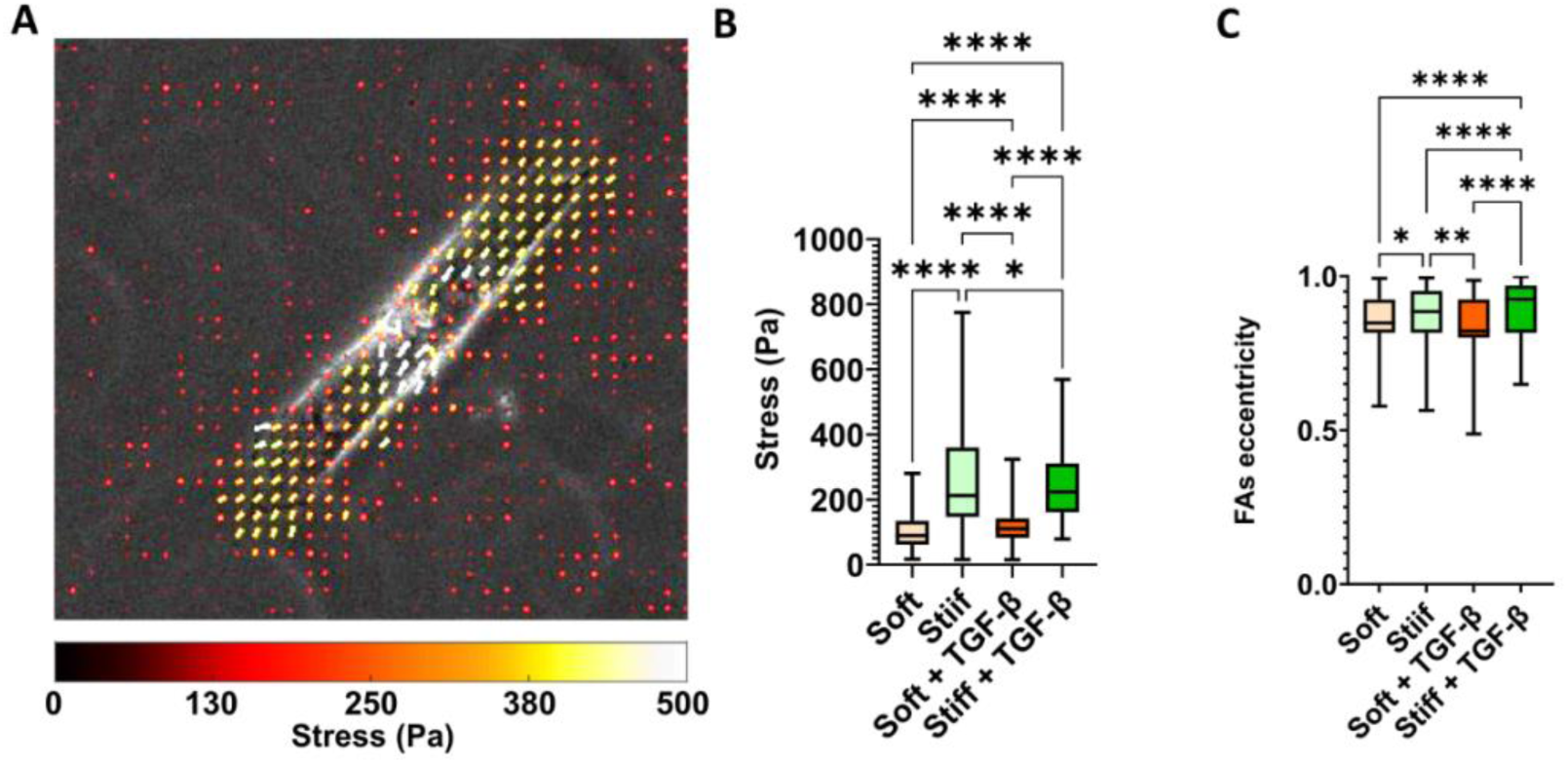
Traction forces exerted by fibroblasts is correlated to substrate stiffness and FA elongation. **A)** Representative overlay of traction map (traction vectors indicated by arrows, where the color indicates the magnitude) on phase-contrast image of fibroblasts cultured for 96 hours on stiff hydrogels without TGF-β treatment. **B)** Mean traction forces applied by fibroblasts on stiff and soft hydrogel with and without TGF-β. Mean of the forces obtained from >10 FOV containing multiple cells on n = 3 substrates for each condition. **C)** Mean FA eccentricity of fibroblasts on hydrogels of different stiffnesses with and without TGF-β treatment. The boxplots represent the median and quartiles of the distributions, with the whiskers indicating the outliers (*p<0.5, **p<0.05, ****p<0.00001).

On soft pAA gels, the measured traction forces was significantly lower (101.3 ± 51.6 Pa) compared to the traction forces exerted by fibroblasts cultured on the stiff hydrogels (270.5 ± 168.6 Pa) (Figure 3B; *p* < 0.0001). This is in agreement with the literature, where it has been shown that forces exerted by fibroblasts are higher when the cytoskeleton are dense with actin fibers[30]. TGF-β induces the incorporation of αSMA into actin fibers after 96 hours (Figure 1B), which is associated with an increase in actin bundle formation and the enhanced contractile potential of fibroblasts[11,28]. Interestingly, we found that TGF-β stimulation evoked different effects on the magnitude of cell traction forces depending on the substrate stiffness. TGF-β-induced cells on the soft gels exerted higher traction forces (121.2 ± 58.1 Pa) compared to the control (*p* < 0.0001), as expected. However, fibroblasts cultured with TGF-β on stiff gels showed slightly lower mean traction forces levels (245.9 ± 103.6 Pa) than the control (*p* = 0.036). The observed increase and decrease in traction forces exerted by fibroblasts on soft and stiff hydrogels, respectively, may be attributed to the stabilization of actin bundles due to changes in FAs turnover, which could lead to less dynamic force generation despite the increased contractility[43,44].

The force exerted by fibroblasts is produced by actomyosin and transmitted through FAs. These FAs play a crucial role in stabilizing actin bundles through a process known as turnover[43,44]. To exert higher forces on substrates, fibroblasts require the formation of mature FAs and a dense network of F-actin bundles[45–47]. This interconnected system ensures the efficient transmission of mechanical forces and supports the cells’ ability to exert substantial traction on their environment. Since the force exerted by the fibroblasts is transferred to the hydrogels through the FAs, we investigated whether the FA morphology present any correlation with the force exerted. We quantified the FAs eccentricity of fibroblasts cultured on soft and stiff hydrogels for 96 hours as a response to the pulling force of the cytoskeleton. FAs were rounder on softer substrates and more elongated on stiffer hydrogels (Figure 3C; *p* = 0.0418). Moreover, the introduction of TGF-β did not cause FAs to adopt a more elongated shape on soft hydrogels (Figure 3C). However, the addition of TGF-β was correlated with higher traction forces exerted by the cells compared to the control (Figure 3C). In contrast, on stiff hydrogels, TGF-β induced a more elongated FA shape (Figure 3C; *p* < 0.0001), even though it resulted in slightly lower force (Figure 3B). This is consistent with a previous report that myosin II-mediated intracellular tension is involved in the regulation of FAs stability and elongation, but without affecting their growth rate[48].

These results confirm that, although TGF-β promotes the formation of more stable FAs, it is insufficient to drive fibroblasts toward a fully contractile myofibroblast phenotype on a soft substrate. On stiff substrates, the TGF-β-induced enhanced stability of FAs leads to slightly increased cell traction forces and FA elongation, whereas on soft substrates, fibroblast activation via TGF-β is attenuated by the low substrate stiffness, which impairs the stability and maturation of focal adhesions. Consequently, only a marginal increase in traction force is generated on the substrate, consistent with limited FA stabilization. This weak mechanical feedback affects stress fiber assembly and is insufficient to induce full differentiation of fibroblasts into myofibroblasts. Overall, these findings suggest that cell migration is regulated by FA turnover, which is largely dictated by environmental cues such as substrate stiffness.

### Stiffness sensing regulates fibroblast migratory behavior

On soft hydrogels, fibroblasts maintain a rounded morphology (Figure 4A). This is in line with the impaired FA-mediated contractility (Figure 3B). As expected from the literature[45,46], when cells are unable to form F-actin stress fibers, they exert weaker pulling forces on the substrate, resulting in fewer and smaller adhesion events, such as on our soft substrates (Figure 1C). In contrast, TGF-β-treated cells displayed a less rounded morphology (Figure 1, Figure 4A), likely due to their ability to exert greater traction forces on the substrate. On stiff hydrogels, fibroblasts typically adopt a spindle-shaped morphology under normal conditions (Figure 4A). When treated with TGF-β, fibroblasts on stiff hydrogels exhibited larger FAs (Figure 4B; *p* < 0.0001). On stiffer substrates, fibroblasts were able to form mature FAs (Figure 1D), which occur through strong cell–substrate interactions and substrate resistance to actomyosin pulling forces. Although TGF-β treatment induced changes in FA size, resulting in larger FAs on stiff hydrogels (Figure 3C), no significant alterations in FA size were observed on soft hydrogels, even following TGF-β-induced activation (*p* > 0.9).

**Figure 4.**
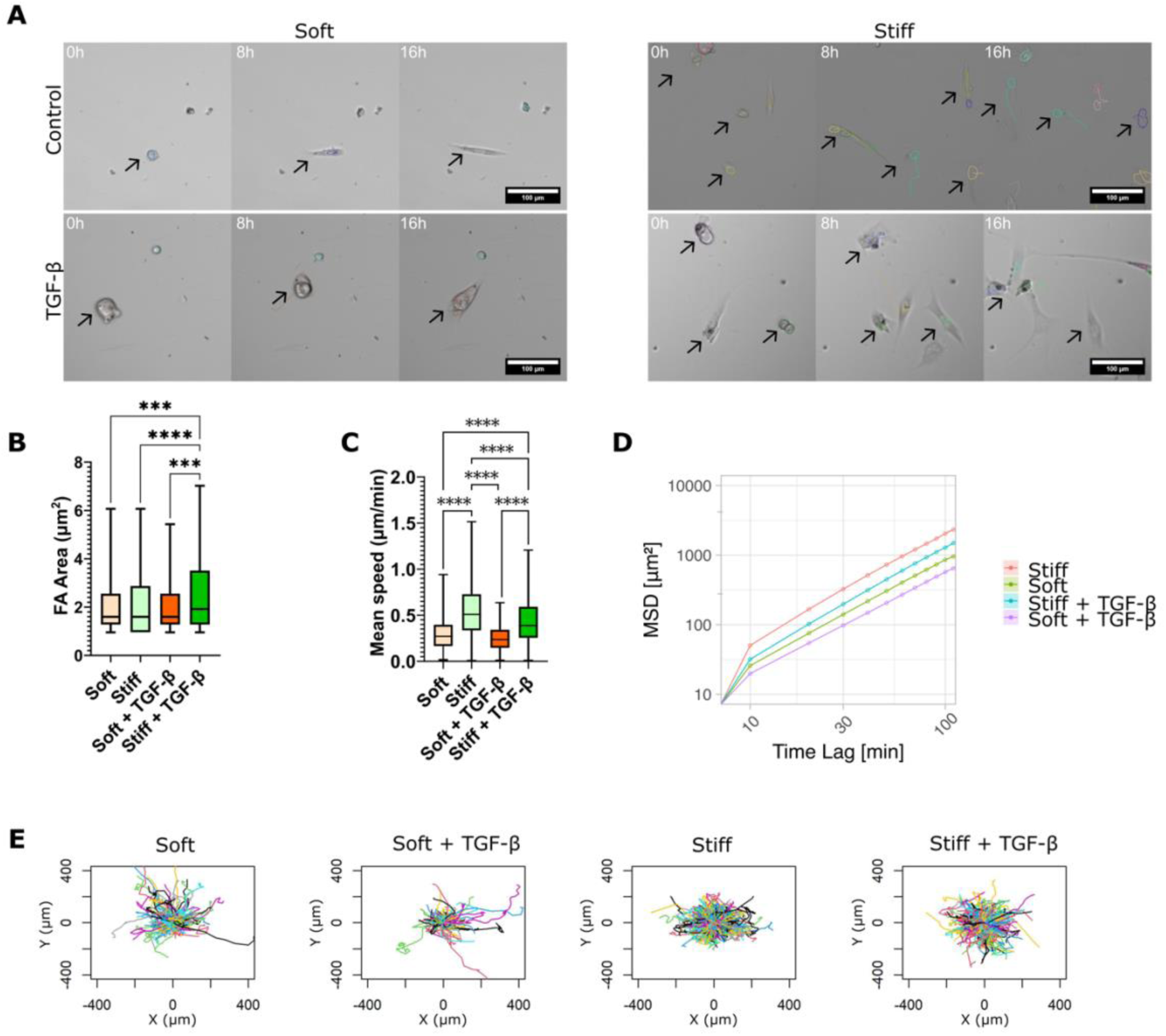
Fibroblast migration is influenced by stiffness sensing. **A)** Fibroblasts cultured on hydrogels with different stiffnesses (soft = 9 kPa; stiff = 30 kPa) for 96 hours with and without TGF-β treatment and tracked (black arrows) over time for 16 hours. **B)** Mean FA area of fibroblasts cultured on hydrogels with and without TGF-β treatment at different stiffnesses. **C)** Mean speed of fibroblasts cultured on hydrogels presenting different stiffnesses with and without TGF-β treatment. Data in panel B and C are shown as the boxplots representing the median and quartiles of the distributions, with the whiskers indicating the outliers (***p < 0.005, ****p < 0.00001). **D)** MSD obtained from single cell migration analysis of fibroblasts cultured on soft and stiff hydrogels in the presence or absence of TGF-β for 96 hours, plotted as mean ± SD. **E)** Trajectories of nuclei tracking of fibroblasts cultured on soft and stiff hydrogels in the presence or absence of TGF-β.

The pulling force generated by the actomyosin and the consequent FA maturation are essential for cell migration[25] We have already observed in the previous sections that, when activated by TGF-β, fibroblast increased the actin stress fibers formation (Figure 1) and gained a higher motility (Figure 2C). To understand if the stiffness mechanosensing influences fibroblasts migration independently to FMT, we quantified the migratory behavior of fibroblasts on substrates of different stiffnesses, both in the presence and absence of TGF-β (Fig. 4C). Fibroblasts cultured for 48 hours on soft and stiff hydrogels showed a clear influence of stiffness on their mean speed. Fibroblasts exhibited a migration speed of 0.31 ± 0.20 μm/min on soft hydrogels, significantly lower than 0.39 ± 0.19 μm/min on stiff hydrogels (Figure S3A; *p* < 0.0001). Similarly, fibroblasts cultured for 96 hours on stiff hydrogels exhibited higher mean speeds (0.56 ± 0.29 μm/min) compared to those cultured on soft hydrogels (0.32 ± 0.22 μm/min) (Figure 4C; *p* <0.0001). On both stiffnesses, when the cells were treated with TGF-β, their mean speed decreased compared to that of fibroblasts cultured without TGF-β, with the difference being less pronounced on softer substrates (Figure 4C; *p* = 0.51 on stiff and *p* < 0.0001 on soft substrates). Interestingly, analysis of the mean squared displacement (MSD) of fibroblasts on both soft and stiff hydrogels revealed that TGF-β treatment resulted in reduced migration over time, regardless of substrate stiffness (Figure 4D and 4E).

After 96 hours in the presence of TGF-β, fibroblast motility was notably affected by decreasing fibroblasts mean speed by FA stabilization, but this effect was only prominent on stiff substrates, where fibroblasts transitioned into myofibroblasts. This transition was associated with reduced motility and the formation of larger FAs. These findings underscore the critical role of substrate stiffness in mechanosensing and its regulation of fibroblast migratory behavior.

## Discussion

Fibroblasts play a crucial role in maintaining and (re)establishing tissue homeostasis. Normally, they exist in a quiescent state with a spindle-shaped morphology, producing and modifying ECM components to maintain tissue structure and architecture. In response to stress, such as injury, wound, or inflammation, fibroblasts are activated, typically due to elevated TGF-β levels. Persistent exposure to activation factors leads to fibroblast differentiation into myofibroblasts via fibroblast-to-myofibroblast transition (FMT). Subsequently, after their activation fibroblasts migrate towards the stress site. This differentiation is characterized by increased contractility, which is enabled by the incorporation of αSMA into the actin cytoskeleton. In this process, not only are fibroblasts exposed to tissue microenvironments with varying stiffness, but the FMT itself also results in modifications of the tissue stiffness. In this study, we asked whether the stiffness of the microenvironment also influence fibroblast phenotype and behavior during FMT and whether the regulation of FMT is guided by mechanosensing machinery independently of biochemical induction with TGF-β.

We first set out to establish the temporal progression and timeline of FMT, as well as the observable cellular characteristics in each stage of FMT. After 48 hours of activation by TGF-β, the F-actin bundles formation, and cytoplasmic αSMA expression were enhanced, whereas after 96 hours, αSMA became incorporated into the F-actin stress fibers. These are consistent with the known phenotypical transition of fibroblasts during FMT, first into proto-myofibroblasts, and afterwards into myofibroblasts[21,29,39,49]. Furthermore, quantitative analysis of cell migration trajectories revealed a clear dependency on the time point analyzed. At the early time point (48 hours), TGF-β-induced cells at the proto-myofibroblast stage showed higher migration speed compared to the quiescent fibroblasts. In contrast, at the later time point (96 hours), the TGF-β-induced myofibroblasts were slower than the quiescent fibroblasts. This phenotype-dependent switch in migration characteristics may have a physiological advantage. Upon activation, fibroblasts should quickly migrate toward the site of tissue insult. This fast migration is facilitated by cytoskeletal rearrangement and the expression of actomyosin bundles, allowing generation of cellular traction forces that are important for cell locomotion[25,50]. After the cells reach the point of interest, proto-myofibroblasts complete the phenotypical change into myofibroblasts, typified by the incorporation of αSMA in the actin bundles. This strongly increases cell contractility and leads to a more stationary behavior[51–53], while at the same time allowing myofibroblasts to remodel the microenvironment.

To mimic the environment stiffness during pathological conditions of tissue, we cultured fibroblasts on polyacrylamide (pAA) hydrogels after 96 hours with Young’s moduli representing healthy (∼9 kPa) and pathological (∼30 kPa) tissue conditions. Fibroblasts cultured for 96 hours on softer hydrogels developed smaller FAs, even when induced by TGF-β. This is in line with previous studies that have shown that cells on soft substrates are unable to produce mature FAs nor form F-actin bundles[7,8]. Conversely, fibroblasts cultured on stiffer hydrogels developed larger FAs when induced by TGF-β. These cells not only produced a dense network of F-actin stress fibers, as observed in the control condition but also incorporated αSMA, indicating a phenotype transition into myofibroblasts which is in agreement with previous knowledge. However, our data suggest that the fibroblast phenotype during FMT is primarily mechanically (rather than biochemically) conditioned by the stiffness of the microenvironment. Specifically, fibroblasts require a stiff environment to form mature FAs and actin stress fibers to be activated, and the addition of TGF-β further promotes the incorporation of αSMA into the actin cytoskeleton to become myofibroblasts.

To further probe the impact of stiffness on FMT, we characterized the migratory behavior of fibroblasts at different phenotypical stages on soft and stiff hydrogels. This is especially pertinent, as the same cytoskeletal components that cells use to sense substrate stiffness are also important for cell migration, and we have already seen that both stiffness and TGF-β lead to significant rearrangements of the actin cytoskeleton and FAs. Additionally, the migration of fibroblasts to the source of tissue stress is a critical first step in tissue repair and regeneration. We found that fibroblasts were more actively migrating and had a higher mean migration speed on stiffer substrates than on softer substrates. This is consistent with the observation that stiffness sensing allows fibroblasts to form a stronger network of actin bundles on stiff substrates, which is needed for cell locomotion[25,29]. Further, we investigated whether changes in the fibroblast migratory behavior were due to the substrate stiffness or to changes in the activation state of the cells. On softer substrates, fibroblasts activated by TGF-β did not exhibit differences in migration compared to the control fibroblasts. However, on stiffer substrates, TGF-β-induced activation significantly decreases the mean speed of the fibroblasts compared to the untreated condition. Soft hydrogels help maintain fibroblasts in a quiescent-like state, with only a slight increase in migration observed when TGF-β is added. In contrast, increasing substrate stiffness alone significantly enhances cell migration and activates fibroblasts. However, while substrate stiffness can drive activation, the presence of TGF-β and their incorporation into the actin cytoskeleton are still necessary for fibroblasts to fully transition into the myofibroblastic phenotype. When fibroblasts are recruited to the site of tissue stress, their migratory speed should increase. To enhance cell motility, fibroblasts must form robust F-actin bundles, which play a crucial role in initiating fibroblast activation and promoting efficient migration[25]. Interestingly, on stiff hydrogels, TGF-β-induced activation leads to a decrease in migration speed, likely because the cells are transitioning into myofibroblasts. This slower migration is consistent with the myofibroblasts’ primary function, which shifts from migration to microenvironment remodeling once they reach the site of tissue stress.

To further understand how substrate stiffness influences FMT and cell–matrix interactions, we performed traction force microscopy (TFM) analysis. Previous studies have demonstrated that fibroblasts on stiffer hydrogels tend to form robust F-actin bundles, while those on softer hydrogels do not. This leads to greater forces being exerted on stiffer substrates due to enhanced actomyosin pulling forces, whereas softer substrates experience lower forces. Interestingly, when fibroblasts were treated with TGF-β, we observed an increase in the force exerted on softer substrates, which suggests that TGF-β enhances the mechanosensitivity of fibroblasts even in less stiff environments. Conversely, on stiffer substrates, TGF-β treatment led to a decrease in force exertion. This finding aligns with our earlier observation that TGF-β-induced activation on stiff substrates results in a transition to the myofibroblast phenotype, which is characterized by reduced migration speed and increased focus on matrix remodeling rather than force generation for migration.

These results highlight substrate stiffness, stiffness sensing, and stiffness response, as a key determinant of fibroblast phenotype. A recent computational study indicated that the spatial and morphological properties of FAs determine the stiffness perceived by cells[54]. As such, we further explored the relationship between cell migration, FA size, and substrate stiffness, as well as how these factors influence the phenotypical transition during FMT. We found that on soft hydrogels, fibroblasts maintained smaller FAs and did not undergo significant phenotypical changes, even when treated with TGF-β. In contrast, on stiff hydrogels, fibroblasts developed larger FAs when induced by TGF-β, leading to the formation of dense F-actin stress fibers and the incorporation of αSMA, indicative of the myofibroblast phenotype. This suggests that substrate stiffness plays a critical role in regulating the fibroblast phenotype during FMT, with TGF-β serving as a necessary biochemical signal to drive the full transition to myofibroblasts (Figure 5). Importantly, the reduced migration speed observed on stiff substrates following TGF-β treatment reflects the functional shift of myofibroblasts from migratory cells to agents of microenvironmental remodeling, underscoring the complex interplay between mechanical and biochemical cues in tissue repair and fibrosis.

**Figure 5.**
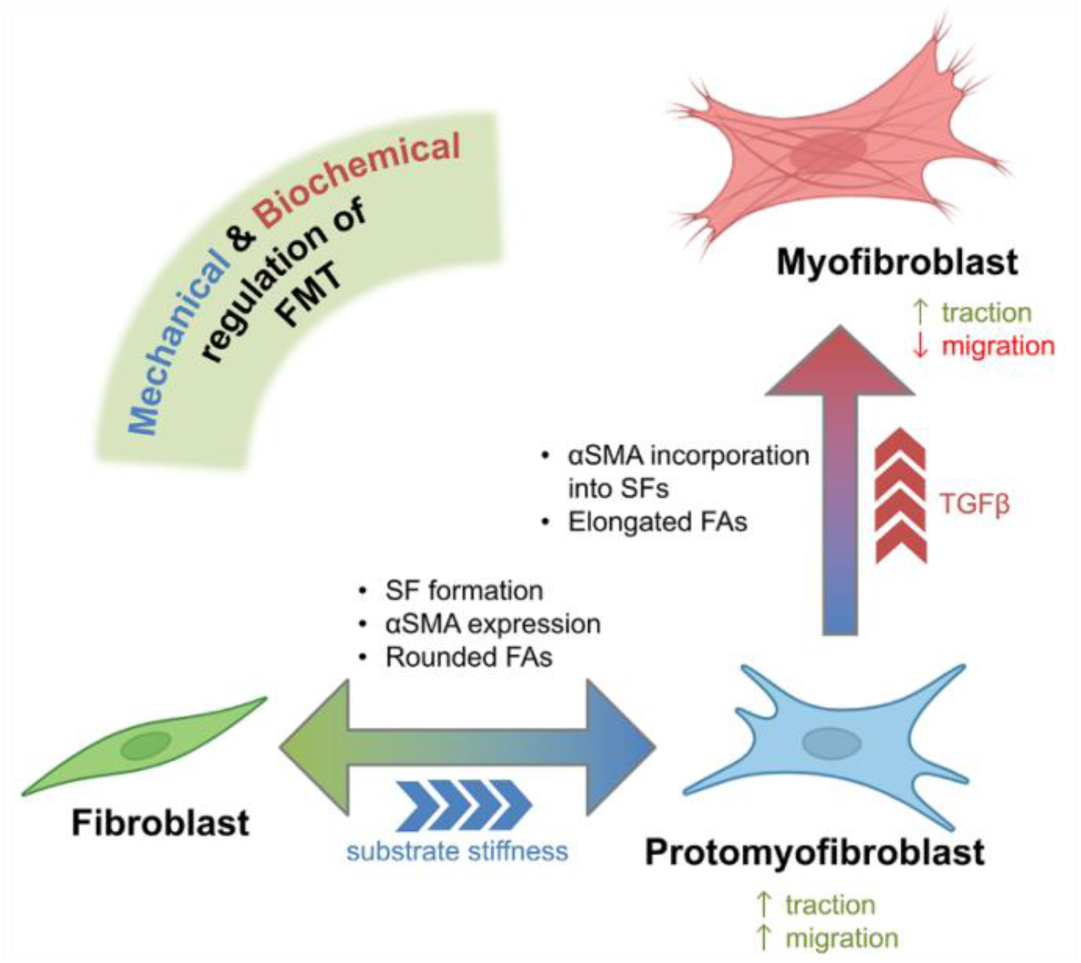
A working model of fibroblast activation and migration dynamics during FMT in the presence of mechanical and biochemical cues in the tissue environment. Fibroblasts sense substrate stiffness and progressively form mature focal adhesions (FAs) and actin stress fibers as stiffness increases. Exposure to TGF-β in these mechanically activated setting allows incorporation of αSMA into actin stress fibers, leading to myofibroblast differentiation. Migratory behavior follows a biphasic pattern: fibroblast speed initially increases on stiffer substrates, then decreases upon myofibroblast transition, demarcating a shift from active motility to matrix remodeling.

## Conclusion

Our study provides insights into the multifaceted regulation of fibroblast behavior by integrating both mechanical and biochemical factors, which are essential for understanding critical biological processes such as wound healing. We uncover a novel and intricate crosstalk between stiffness mechanosensing and fibroblast phenotypical development that plays a decisive role in regulating fibroblast migratory behavior. Moreover, we demonstrate that stiffness, when coupled with phenotypical changes, not only modulates migratory behavior but also results in significant differences in cell contractility. More broadly, this study introduces a novel perspective on the complex interplay between morphological, mechanical, and phenotypic characteristics of fibroblasts. We show that alterations in any one of these aspects can evoke substantial changes in the others. This may be useful for developing more effective strategies in wound healing and tissue regeneration, such as by manipulating the microenvironmental stiffness and its sensing as well as the cellular morphological and mechanical behavior.

## Materials and Methods

### Cell Culture

Normal Human Dermal Fibroblasts (nhDFs, Lonza, CC-2511) were cultured in Dulbecco’s Modified Eagle Medium (DMEM; Thermo Fisher, 41966-029) supplemented with 10% fetal bovine serum (FBS; Biochrom AG) and 1% penicillin/streptomycin (P/S; Biochrom AG). The culture medium was refreshed with DMEM containing 3% FBS and 1% P/S one day before the start of the experiment. Cells were seeded at a density of 471 cells/cm² onto 13 mm diameter patterned glass coverslips or pAA gels in DMEM with 3% FBS and 1% P/S. To ensure localized cell attachment, 200 µl of the cell suspension was carefully pipetted onto the glass surface or gel and incubated at 37°C for 30 minutes. Once cell attachment was observed, the wells were filled with additional medium to wash away non-adherent cells. The nhDF cultures were then incubated under the specified experimental conditions at 37°C in a cell-culture incubator with 5% CO_2_.

### pAA hydrogels

Hydrogels were prepared using a mixture of 40% acrylamide, 2% bis-acrylamide, ammonium persulfate (Bio-Rad), and tetramethylethylenediamine (TEMED; Bio-Rad) in phosphate-buffered saline (PBS). A 12 µL droplet of this solution was pipetted onto bind-silane functionalized glass slides, followed by the placement of a 13 mm diameter coverslip on top. The hydrogels were allowed to polymerize for 60 minutes at room temperature, after which PBS was added and the gels were incubated for an additional hour. Subsequently, the coverslip was carefully removed, and the gels were thoroughly washed with PBS (Sigma-Aldrich). The hydrogels were stored in PBS at 4°C until further use. For the TFM analysis, 200-nm-diameter dark-red fluorescent carboxylate-modified beads (F8807, ThermoFisher) were added at a concentration of 1% to the pAA gel solution.

### Fluorescence staining and imaging

After the experiments concluded at 48 and 96 hours, cells were fixed with 3.7% paraformaldehyde for 20 minutes at room temperature, followed by a 10-minute treatment with Triton X-100 to precipitate cytoplasmic proteins. Staining was performed using 4’,6-diamidino-2-phenylindole dihydrochloride (DAPI; Sigma-Aldrich, D9542), vinculin (mouse anti-vinculin IgG1 antibody, 1:600; Sigma, V9131), F-actin (Phalloidin-Atto647), fibroblast activation protein (FAP; rabbit polyclonal anti-FAP, 1:600; Sigma-Aldrich), and α-smooth muscle actin (αSMA; mouse anti-αSMA IgG2A, 1:600; Sigma-Aldrich). Additionally, Atto 590-conjugated Phalloidin (Merck Life Science NV) was used for further staining. The secondary antibodies employed were Alexa Fluor 488-conjugated goat anti-mouse IgG2A for αSMA, Alexa Fluor 647-conjugated goat anti-mouse IgG1 for vinculin, and Alexa Fluor 647-conjugated goat anti-rabbit for FAP (all from Molecular Probes). Fluorescence images were acquired using a Leica SP8X confocal laser scanning microscope equipped with 20×/0.75 or 40×/0.95 objective lenses, operating at a scanning speed of 100 Hz and utilizing gating to enhance the signal-to-noise ratio.

For live cell imaging, fibroblast nuclei were stained with NucBlue (Fisher Scientific, 12303553) by adding one droplet to the culture medium after 48 or 96 hours of culture, followed by a 15-minute incubation. The medium was then refreshed, and the samples were imaged continuously for 16 hours at 37°C in an environment with 5% CO2. Imaging was performed at a frame rate of one frame every 10 minutes, capturing both brightfield (BF) and DAPI signals. Images were acquired using a Leica DMi8 widefield microscope equipped with an HC PL Fluotar 10x / 0.32 objective lens.

### Single-cell migration analysis

Single-cell migration parameters were automatically quantified as described by Maiuri et al41. Cells were seeded on glass or polyacrylamide (pAA) hydrogels and cultured for 48 or 96 hours. NucBlue™ Live ReadyProbes™ Reagent (Hoechst 33342, Invitrogen) was added for 15 minutes, followed by medium refreshment. Cell displacements were monitored every 10 minutes using fluorescence and phase-contrast microscopy over 16 hours period at 37°C in an environment with 5% CO_2_. Imaging was conducted on a humidity- and temperature-controlled inverted wide-field microscope (Leica DMi8) equipped with a 20x/0.4 HC PL Fluotar objective, within an environmental chamber.

Images were processed using the BioFormats plugin, with channels separated for each position. Nuclei were tracked automatically using a custom macro in CellTracker. The resulting trajectories were used as input for a custom R-based analysis to compute various migration parameters. Mean cell speed was calculated by averaging displacements between consecutive frames over time, while the mean square displacement (MSD) quantified the deviation of a cell’s position from a reference point over time. For each cell migration analysis we merged the trajectories of each biological replicates where for each experiment we have 3 technical replicates and >3 biological replicates with >300 cells per condition.

### Traction Force Microscopy

Cells and fluorescent beads were imaged overnight for 16 hours using Nikon Eclipse Ti2-LSM with 20× magnification at 37°C in an environment with 5% CO_2_. The automated analysis was carried out using a handmade MatLab code. The reference image was obtained after removal of the cells. Timelapse images were aligned and cropped according to the reference image. Bead displacements were computed using Particle Image Velocimetry. Cell tractions were computed by Fourier transform-based traction microscopy. Traction force magnitude was calculated as median value over space (individual cell) followed by mean value over time[55].

### Automated image analysis of cell and FA morphology

The automated image analysis workflow comprised three sequentially automated steps: image preprocessing (using Python), morphology analysis (using CellProfiler), and data postprocessing (using Python). Initially, the image collection, stored in native .lif format, was processed using a custom Python library. The images were converted into single .tiff files, which were then organized into separate folders, with each folder containing individual channel images and compressed stacks generated through the maximum projection function. The final preprocessing step involved adjusting the image saturation to 0.35 by modifying the contrast and applying a Top Hat filter to the focal adhesion (FA) channel images. These preprocessed images were then used as inputs for the CellProfiler pipeline.

In CellProfiler, nuclei were first identified using the IdentifyPrimaryObjects tool, where grayscale images were segmented using the Otsu two-class method, with an option for manual refinement via the EditObjectsManually tool. Cell boundaries were defined using the IdentifySecondaryObjects tool and filtered similarly to the nuclei. For the FA identification, cell boundaries were initially masked, followed by conversion into analyzable objects and segmentation via the thresholding function. Quantitative measurements, such as object size and shape, as well as area occupied by the image, were obtained using tools like MeasureObjectSizeShape and MeasureImageAreaOccupied. The resulting data were saved as .xlsx files.

Postprocessing of the analyzed data (.xlsx files) related to specific experimental conditions was performed using another custom Python library. This library processed the data to extract desired parameters per object and saved the results as separate .xlsx files, while also generating summary graphs for individual objects and comparative graphs across different experimental conditions.

### Statistical analysis

Data are plotted either as their full distribution (using boxplots) or descriptively (using mean ± SD)in, as indicated in the respective figure captions. Statistical analyses were performed using GraphPad Prism 9. One-way ANOVA was used to analyze statistical significance of different experimental conditions, including the control condition. Furthermore, we used Mann–Whitney test multiple comparison test to compare the mean of each group with the control condition.

### Resource availability Lead contact

Further information and requests for resources and reagents should be directed to and will be fulfilled by the lead contact, Nicholas A. Kurniawan.

### Materials availability

This study did not generate new unique reagents.

### Data and code availability

The data supporting this study’s findings are available from the lead contact upon reasonable request.

## Acknowledgments

The research project has received financial support from the European Research Council (ERC) under the European Union’s Horizon 2020 research and innovation program (CoEvolve, grant no. 851960 for N.A.K.), and from the Dutch Research Council (NWO; the Gravitation Program “Materials Driven Regeneration”, grant no. 024.003.013 for C.V.C.B and N.A.K.). The authors thank Adria Villacrosa Ribas and Leon L.H. Hermans from the Soft Tissue Engineering and Mechanobiology group for insightful discussions. Microscopy was performed at the LCTE Microscopy Facility, TU/e.

## Author contribution

MD: conceptualization, methodology, validation, formal analysis, investigation, data curation, writing – original draft, writing – review & editing, visualization; PvdB: validation, formal analysis, investigation; SP: formal analysis, data curation, writing – review & editing; PM: conceptualization, software, formal analysis, writing – review & editing; CVCB: resources, writing – review & editing, supervision, project administration; NAK: conceptualization, resources, write original draft, writing – original draft, writing – review & editing, supervision, project administration, funding acquisition.

## Declaration of interests

The authors declare no competing interests.

## Notes

### Competing Interest Statement

The authors have declared no competing interest.

