## Supplementary figures for "Substrate stiffness modulates phenotype-dependent fibroblast contractility and migration independent of TGF-β stimulation"

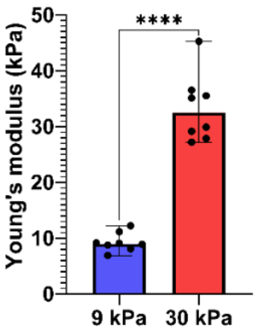


**Figure S1**. Young's modulus of pAA hydrogels. ***** p<0.0001(Unpaired t test)

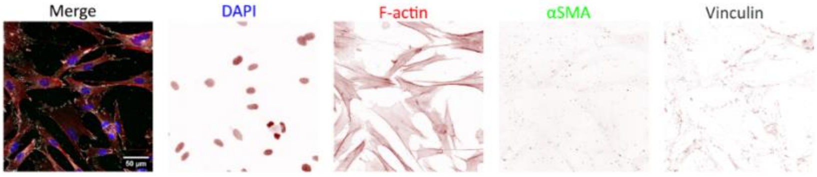


**Figure S2**. Fibroblasts cultured 96 hours on glass substrate in presence of AIIB2 were stained for DAPI (blue), αSMA (green), f-actin (red) and vinculin (grey). Scale bar = 50 μm

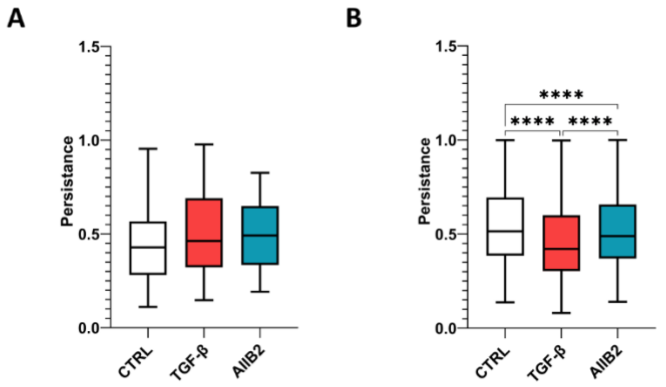


**Figure S3**. **Fibroblast persistence shows differences in different phenotypical states**. **A-B)** Persistence of fibroblasts cultured on glass for (A) 48 hours and (B) 96 hours. **** p<0.0001 (One-way ANOVA)

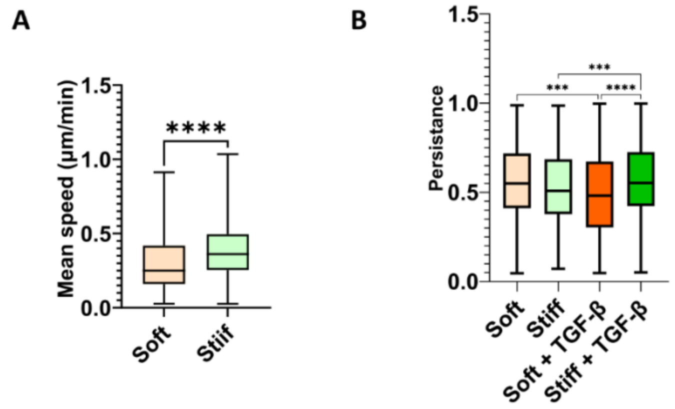


**Figure S4. A)** Mean speed of fibroblasts cultured on soft and stiff hydrogels for 48 hours. **B)** Persistence of fibroblast cultured on pAA hydrogels for 96 hours. ***p<0.005 , ****p<0.0001 (Unpaired t-test and One-way ANOVA)
